# A Reduced Mechanobiological Framework for Platelet Priming: From Hemodynamic Shear to Mechanosensitive Calcium Entry

**DOI:** 10.64898/2026.08.03.742655

**Authors:** Yuxin Chen, Xiaochen Liu, Daniele Vigolo, Mackenzie Siyuan Zhuang-Hall, Ken-Tye Yong

## Abstract

**Background:** Platelet activation in flowing blood is a multiscale process in which vessel-scale hemodynamics, red blood cell (RBC) mechanics, adhesive receptor interactions, and intracellular signalling jointly determine thrombotic risk. Individual components are well studied, but a single reduced description that carries each explicitly from vessel-scale flow to mechanosensitive calcium entry, with dimensionally consistent couplings, remains uncommon.

**Objectives:** We develop and analyse a reduced, six-module mechanobiological framework for platelet priming spanning the cascade from hemodynamic shear to mechanosensitive calcium entry, and we delineate which elements are supported by existing evidence and which are new, testable hypotheses.

**Methods:** The framework comprises six coupled modules: (I) hemodynamic forcing from the incompressible Navier–Stokes equations, with an objective principal-strain-rate measure for extensional flow; (II) RBC-mediated platelet margination and near-wall delivery, closed by a near-wall arrival flux; (III) von Willebrand factor (VWF) activation with a bounded kernel and glycoprotein Ib*α* (GPIb*α*) catch-slip capture, resolved through an explicit contact area and a bond-dependent mobility that progressively immobilises wall-interacting platelets; (IV) a single-load membrane-stimulus formulation; (V) mechanosensitive gating and a dimensionally consistent cytosol–store calcium model with extracellular influx; and (VI) a phenomenological mechanical-memory state. We formally derive that the single-platelet stochastic dynamics and the continuum population balance form a Fokker–Planck pair, with the spatially varying diffusivity handled by an explicit drift correction.

**Results:** The framework yields a family of mechanochemical dimensionless groups delineating priming regimes. Its central prediction is reformulated as a falsifiable, history-sensitive signature: in a conditioning–test protocol, a low-tension conditioning block charges the memory state, and a fixed sub-threshold test pulse then reports a delay-dependent calcium facilitation that decays on the memory time *τ*_*m*_ and is distinguishable from no-memory gating, channel adaptation, and residual-calcium priming. We show explicitly that the previously proposed pulsatile-versus-monotone contrast is a nonlinear convexity/thresholding effect of the gating nonlinearity—its difference-in-differences is approximately zero— and is therefore *not* a valid test of memory; the conditioning–test signature is. A second prediction links RBC stiffening to reduced near-wall delivery and captured-platelet calcium response, upstream of intrinsic platelet signalling.

**Conclusions:** The framework provides a dimensionally consistent, mechanistically grounded and hypothesis-generating description linking hemodynamic forcing to mechanosensitive calcium entry. It demonstrates how history-dependent platelet priming may arise from a phenomenological sensitisation state and proposes a conditioning–test protocol for comparison against adhesive, channel and intracellular-store persistence. The framework is calibratable rather than validated, and the quantitative outputs shown use representative uncalibrated parameters.

## 1 Introduction

Arterial thrombosis—the pathological occlusion of a vessel by a platelet-rich thrombus— underlies acute myocardial infarction, ischaemic stroke, and peripheral vascular disease. Its initiation is governed not by blood chemistry alone but by hemodynamics: vessel geometry and calibre set local shear-stress distributions, which determine whether platelets are transported toward the wall, whether VWF unfolds and captures flowing platelets, and whether intracellular signalling is mechanically triggered.

This mechanobiological perspective has strong experimental support. Hellums established a critical shear-stress threshold for platelet aggregation, making hemodynamic force a direct activating stimulus [1]. Shear-induced platelet aggregation operates at elevated shear even without classical agonists [2, 3]; VWF unfolding under extensional flow exposes the A1 domain that captures GPIb*α* [4, 5]; and near-wall platelet enrichment is controlled by RBC deformability and hematocrit [6]. Two further nodes were added over the past decade: shear *gradients*, not merely peak shear, drive localised thrombus formation [7]; and Piezo1 was identified as a mechanosensitive cation channel on platelets [8–11], giving a molecular identity to shear-triggered calcium entry. Integrin *α*IIb*β*_3_ transitions through an intermediate-affinity state [12], confirming mechano-chemical coupling throughout the cascade.

Integrated thrombosis models already couple substantial portions of this pathway: fluid-mechanical models resolve hemodynamics but assign platelet activation as a threshold on shear without mechanistic transduction [13]; population-balance models resolve coagulation biochemistry but prescribe activation [14]; and cell-resolved models track RBC and platelet dynamics without intracellular state [15]. What is less common is a single *reduced* description that carries the flow field explicitly through RBC-mediated delivery, VWF–GPIb capture, membrane mechanics, channel gating and intracellular calcium, with dimensionally consistent couplings and a stated boundary between what is supported and what is hypothesis. That is the aim of this paper, the first of a two-part series. The cytosolic calcium concentration *c*(*t*) is the output here and the primary biochemical input of the companion paper, which addresses Rap1-mediated integrin activation, paracrine ADP coupling, and aggregate dynamics.

The framework has three design properties. *First*, inter-module couplings are explicit: each module receives defined inputs and produces defined outputs; the few prescribed inputs (notably the agonist-set IP_3_ concentration and several geometric factors) are named as such. *Second*, it is dimensionally consistent throughout, with every flux and coupling audited for units. *Third*, the single-platelet stochastic dynamics and the continuum population balance are shown to form a Fokker–Planck pair (Section 9). We are explicit that the framework is a *calibratable synthesis*, not a validated model.

## 2 Overview of the Framework

The state of a single platelet at time *t* is ***X***(*t*) = (***x***_*p*_, *z, m, n, c, e, h*), where ***x***_*p*_ is position, *z* ∈ [0, 1] the GPIb*α* capture occupancy, *m* ∈ [0, 1] the mechanical-memory variable, *n* ∈ [0, 1] the channel open fraction, *c* the cytosolic Ca^2+^ concentration, *e* the intracellular-store (dense-tubular-system) Ca^2+^ concentration, and *h* ∈ [0, 1] the IP_3_R inactivation fraction. At the population level a distribution *f* satisfies a population-balance PDE whose deterministic characteristics are the single-platelet ODEs (Section 9).

The modules form a cascade with feedback (Fig. 1). Modules I–V are a forward chain; Module III additionally feeds capture back to motion through a bond-dependent mobility *M* (*z*); Module VI is a history integrator, receiving shear, capture, and membrane load, and returning a sensitising signal *λ*_*m*_*m*(*t*) to Module V (Eq. 19). This feedback topology is the structural origin of Prediction 1 (Section 12).

**Figure 1:**
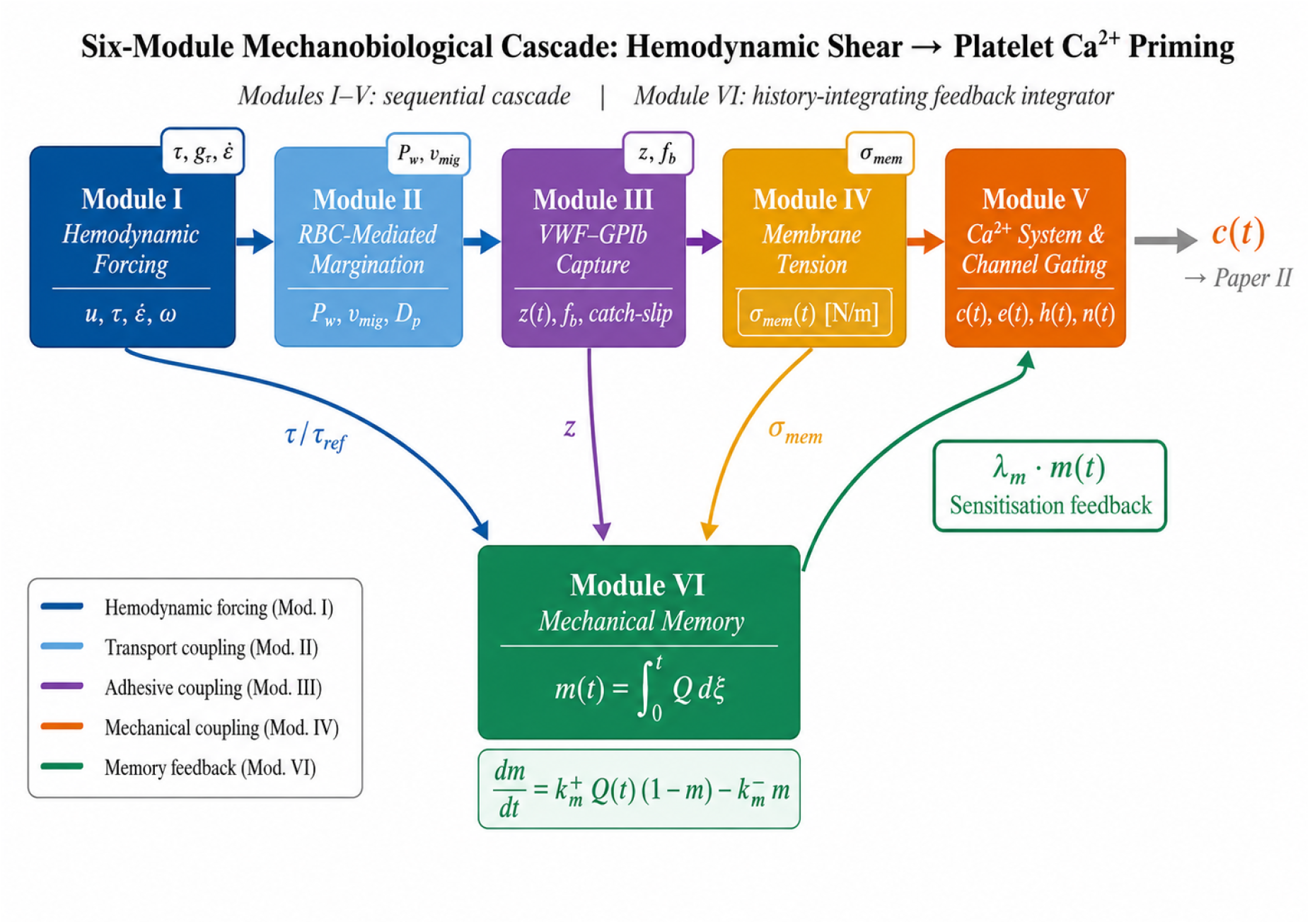
Six-module mechanobiological framework. Modules I–V form the delivery-to-calcium cascade. GPIb occupancy *z* feeds back to platelet motion and diffusion through *M* (*z*) and *D*_eff_(*z*) = *M* (*z*)*D*_*p*_ (labelled *z* → *M* (*z*), *D*_eff_(*z*) → ***x***_*p*_). Module IV supplies the effective mechanosensory load *S*_mech_. Module VI represents a candidate phenomenological history state that modulates mechanosensitive gating through *λ*_*m*_*m*(*t*) (Eq. 19). Cytosolic calcium *c*(*t*) is the output passed to Part II.

## 3 Module I: Hemodynamic Forcing Fields

### 3.1 Governing equations

Blood flow in the vessel domain Ω is modelled as an incompressible viscous fluid,

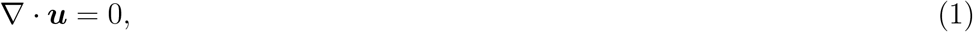

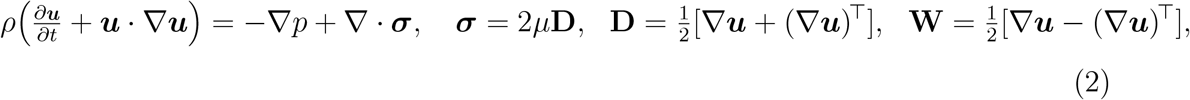

with *ρ* ≈ 1050 kg m^−3^ and *µ* the dynamic viscosity. The Newtonian approximation is appropriate for large-vessel stenotic geometries where shear-driven thrombosis dominates [1, 13]; a Carreau–Yasuda closure may be substituted below 1 mm diameter or mean shear rate 100 s^−1^.

### 3.2 Derived observables

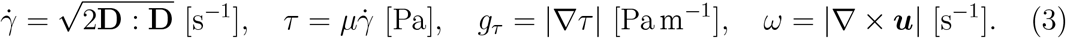

### Objective extensional measure

The extensional forcing on VWF must be frame indifferent. The strain-rate tensor **D** has objective real eigenvalues *λ*_1_ ≥ *λ*_2_ ≥ *λ*_3_ with ∑_*i*_ *λ*_*i*_ = ∇ · ***u*** = 0; we define the extensional rate as the maximum principal strain rate,

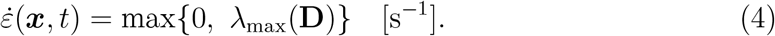

This replaces the earlier scalar 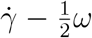, which is not an objective decomposition of extension. We do *not* introduce a separate flow-type parameter: 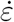 alone enters the VWF kernel (Section 5), and *ω* is retained only as a reported diagnostic.

### 3.3 Platelet trajectory

In the overdamped (Stokes) limit the platelet centre follows the Itô dynamics

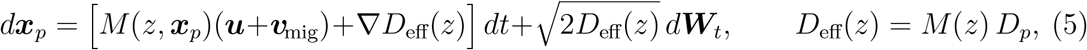

with ***v***_mig_ the RBC-induced margination drift (Module II), *D*_*p*_ the free shear-induced diffusivity (Eq. 7), and ***W***_*t*_ a Wiener process (the drift correction ∇*D*_eff_ is discussed in Section 9). Free platelets have *M* = 1; as GPIb*α* bonds engage near the wall *M* → 0, so both advective mobility *and* diffusivity fall with capture and the platelet is progressively immobilised. A single molecular state *z* therefore controls capture, release, slowing and wall-residence time; there is no separate absorbing-wall mechanism.

## 4 Module II: RBC-Mediated Margination and Near-Wall Delivery

### 4.1 Free-platelet transport

The platelet number density is the marginal *P* (***x***, *t*) = ∫ *f d****q*** of the single distribution *f* over internal states (Section 9); there is no separate free and bound population, because the binding state is carried continuously by *z* and the mobility/diffusivity *M* (*z*), *D*_eff_(*z*) smoothly interpolate between freely advected (*M* = 1) and progressively immobilised (*M* → 0) platelets. Away from the wall, where *M* = 1, the density obeys

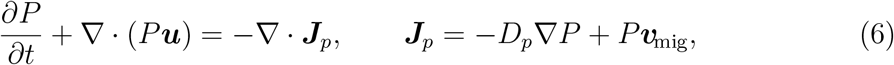

the *M* =1 limit of the population balance (Section 9).

### 4.2 Shear-induced diffusivity and margination drift

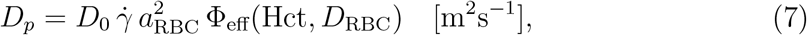

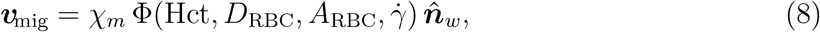

following the shear-diffusion scaling of Eckstein et al. [16], with 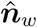 the outward unit normal of the fluid domain constructed from a wall-distance field, and Φ increasing with hematocrit and RBC deformability [6]. Exponents are calibrated in Section 11

### 4.3 Wall-interaction zone and near-wall arrival

Capture is confined to a wall-interaction zone of thickness *δ* ∼ *R*_*p*_,

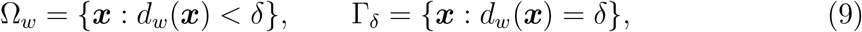

with *d*_*w*_ the wall-distance field and Γ_*δ*_ the outer interface of Ω_*w*_. The physical wall is *reflecting* (no penetration), 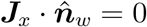, so platelets are not absorbed there; capture is entirely mediated by the molecular state *z* within Ω_*w*_ (Module III). The delivery observable is the near-wall arrival flux across Γ_*δ*_,

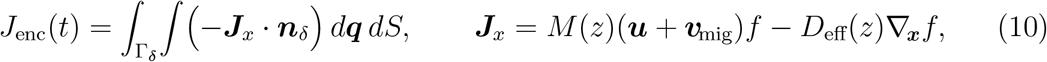

with ***n***_*δ*_ pointing from the wall zone toward the bulk. This replaces the earlier reactive-absorption boundary: there is now a single capture mechanism (*z*), a single population (*f* ), and a conserved platelet number under the reflecting wall.

## 5. Module III: VWF Activation and GPIb*α* Catch-Slip Capture

### 5.1 VWF field and bounded activation kernel

Surface-immobilised VWF *v*_*s*_ ∈ [0, 1] governs capture:

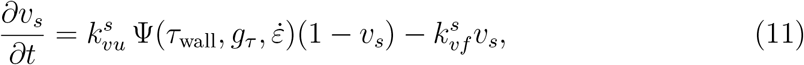

with a *bounded* activation multiplier using saturating Hill factors:

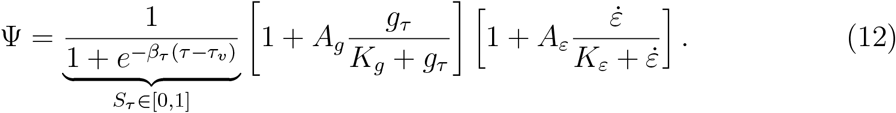

Each bracket saturates to 1 + *A*_*g*_ and 1 + *A*_*ε*_, so Ψ ≤ *S*_*τ*_ (1 + *A*_*g*_)(1 + *A*_*ε*_) is explicitly bounded. The previous unbounded product and the phrase “bounded by calibration” are withdrawn. Activation rises above *τ*_*v*_ ≈ 2–5 Pa [5], is enhanced by shear gradients [7] and by extensional flow [5].

### 5.2 Capture kinetics, bond number, and mobility

Let *z*(*t*) ∈ [0, 1] be the fractional GPIb*α* occupancy. Association requires wall proximity; dissociation does not:

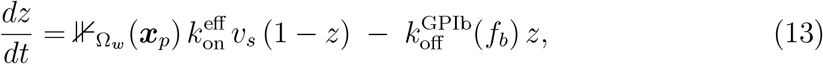

where 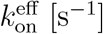 is an effective first-order on-rate (absorbing *R*_GPIb_, *A*_con_, and the 2D association constant). Crucially, the dissociation term is *not* gated by the wall indicator, so a platelet leaving Ω_*w*_ with *z >* 0 correctly relaxes rather than freezing. The number of engaged bonds and total receptor count are

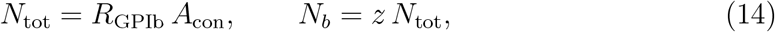

so *N*_*b*_ is the *expected* bond number (a continuous mean, not an integer). The external load *F*_ext_ (Module IV) is shared over the engaged bonds:

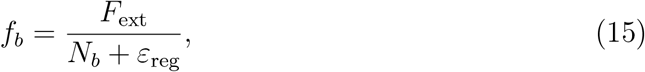

with *ε*_reg_ *>* 0 a stated regulariser keeping *f*_*b*_ finite as *N*_*b*_ → 0; where an integer-resolved bond count is required, Eqs. (13)–(15) should be replaced by a stochastic bond model, and *N*_*b*_ here is its mean-field limit. The bond-dependent mobility entering Eq. (5) is

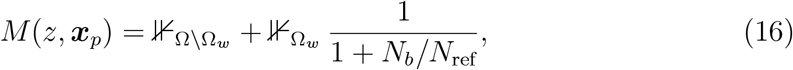

so away from the wall *M* = 1 and within Ω_*w*_ the platelet slows progressively as bonds engage (*M* → 0 only as *N*_*b*_ → ∞); we describe this as progressive slowing, not a discrete arrest switch.

### 5.3 Two-pathway catch-slip dissociation

Unlike the monotonic Bell slip law [17], GPIb*α*–VWF-A1 bonds show a catch regime followed by slip, established by single-molecule force spectroscopy [18, 19]. We use

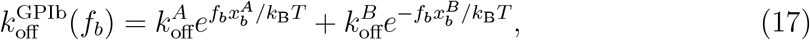

with a lifetime maximum at the catch minimum, consistent with the framework of Evans and Ritchie [20] and directly parameterisable from Yago et al. [18]. Liu et al. [21] support rapid VWF–GPIb–A1 capture of nonactivated platelets; this does not fix the two-exponential parameters or the equal-load-sharing assumption, which remain calibration targets.

## 6 Module IV: Membrane Stimulus from the Transmitted Load

The external load is *F*_ext_ = *F*_shear_ + *F*_coll_ with 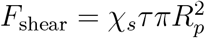 and *F*_coll_ = *χ*_*c*_*f*_coll_⟨*I*_coll_⟩. Engaged bonds *balance* this load (∑_*j*_ ***F***_bond,*j*_ +***F***_hyd_ +***F***_coll_ = **0**), recovering *f*_*b*_ = *F*_ext_/*N*_*b*_ and removing the earlier additive double counting. The mechanically relevant stimulus is the fraction of load transmitted to the cortex,

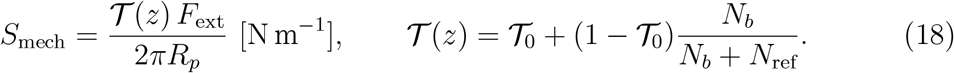

We call *S*_mech_ an *effective mechanosensory load*, not a bilayer tension: it is a lumped, calibrated stimulus, and we do *not* equate it to patch-clamp Piezo1 bilayer-tension thresholds. The normalised stimulus used downstream is 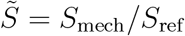, and its half-activation 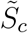 (Eq. 19) is a calibrated parameter fit to platelet mechanosensitive-entry data, not read from single-channel tension measurements.

## 7 Module V: Mechanosensitive Gating and Calcium Dynamics

### 4.1 Biological basis

Shear-mediated Ca^2+^ entry occurs in human platelets and is reduced by gadolinium and by the mechanosensitive-channel blocker GsMTx4 [10]. Piezo1 is gated by bilayer tension [9], and Piezo1 gain-of-function accelerates thrombosis [11]. GsMTx4 acts as a gating-modifier of Piezo1 rather than a clean pore block [22], a point that bears directly on the interpretation of Prediction 1. We treat Piezo1 as the principal mechanically gated entry portal, while allowing effective contributions from other mechanically responsive channels (e.g. TRPC). ATP-gated P2X1 is a ligand-gated channel; it is *not* included in the mechanically gated state *n* and belongs to the agonist-mediated signalling addressed in Part II.

### 7.2 Channel open fraction and the competing-model set

The baseline (memory) model M1 uses a single open fraction with memory sensitisation:

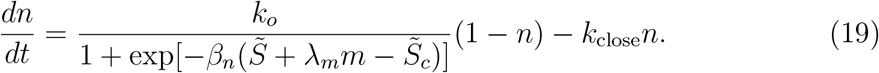

Because a single first-order gate cannot itself distinguish memory from channel adaptation, we define three competitor models sharing the identical calcium readout (Section 12): **M0** (no memory, *λ*_*m*_ = 0); **M2** (channel adaptation, a closed–open–inactivated scheme 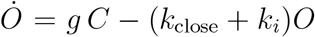, *İ* = *k*_*i*_*O* − *k*_*r*_*I, C* = 1 − *O* − *I*, calcium driven by *O* [23]); and **M3** (residual calcium, M0 with slow cytosolic clearance so baseline Ca^2+^ carries over between stimuli). These are the alternative hypotheses that any claimed memory effect must outperform.

### 7.3 Cytosol–store calcium model with extracellular influx (dimensionally audited)

Following the Li–Rinzel [24] reduction of the De Young–Keizer [25] model, with rapid-buffer effective constants [26]:

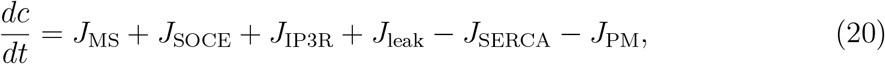

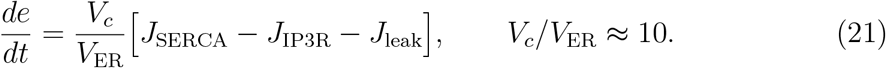

Here *e* denotes the platelet intracellular store (the dense tubular system, an ER-derived membrane system); the Li–Rinzel structure is used as a *reduced store model*, not a literal morphological model of a nucleated-cell ER. Store-operated entry (SOCE) appears *only* in the cytosolic balance; it refills the store only indirectly through SERCA (the Hill exponent satisfies *r* ≥ 1, ensuring the positive-part term 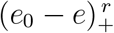 is locally Lipschitz). Every flux is dimensionally consistent [*µ*M s^−1^]: rate constants premultiplying a concentration difference carry [s^−1^], whereas saturating (Hill) fluxes use explicit maximal velocities *V*_•_ [*µ*M s^−1^] with dimensionless gates:

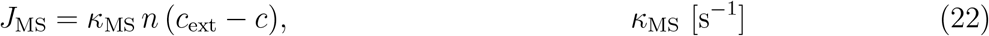

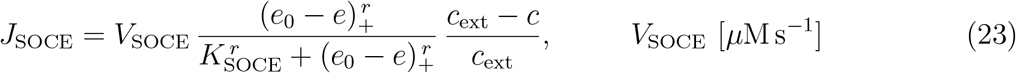

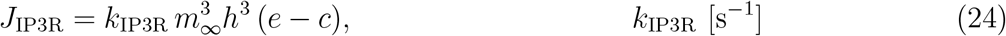

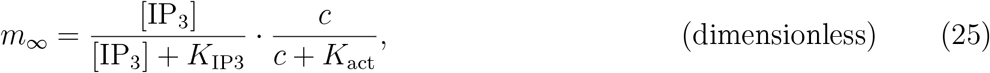

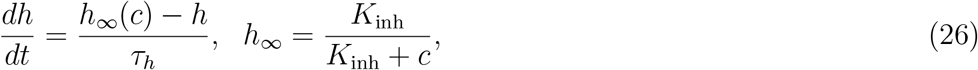

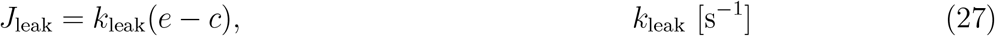

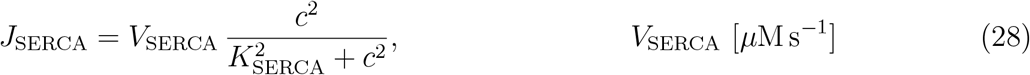

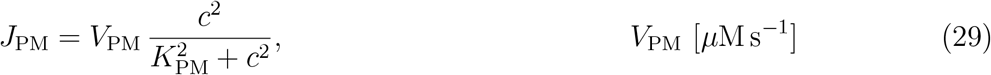

The store leak *J*_leak_ sets the resting balance, and plasma-membrane extrusion *J*_PM_ (PMCA/NCX) is distinct from a generic clearance term. Here *c, e* are free (buffered) concentrations; the IP_3_ concentration [IP_3_] is agonist-set and prescribed in Part I (dynamic in Part II). Store-operated entry follows the physiology of Prakriya and Lewis [27].

## 8 Module VI: Mechanical Memory

We introduce *m*(*t*) ∈ [0, 1]:

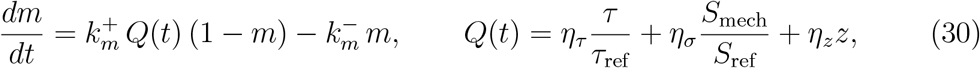

with dimensionless weights *η*_•_ so *Q* is dimensionless. For *m* ≪ 1 this is the convolution 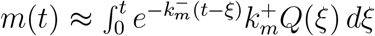, an exponentially weighted average with memory time 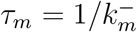.

**Remark 1** (What memory does and does not do). *In the linearised limit the periodicmean memory is* 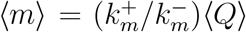, *which depends on the* mean *of Q and not on waveform shape. Consequently equal-mean monotone and pulsatile forcing produce nearly identical mean memory, and the memory model does* not *predict a pulsatile-versus-monotone advantage at equal mean (Section 12); this also holds when the equation is driven by geometry-resolved stenotic shear histories rather than idealised waveforms (Section 12.2). What it does predict is* history dependence at the level of individual stimuli: *a conditioning stimulus transiently sensitises the response to a subsequent test stimulus, decaying on τ*_*m*_. *This conditioning–test facilitation—not a mean-waveform effect—is the history-sensitive, falsifiable signature that the conditioning–test protocol is designed to probe*.

## 9 Single-Platelet Dynamics and the Population Balance

At the single-platelet level spatial motion is stochastic (capture-dependent diffusivity *D*_eff_(*z*) = *M* (*z*)*D*_*p*_) while internal states are deterministic. For a spatially varying diffusivity the Itô SDE requires a drift correction so that the population diffusion is Fickian:

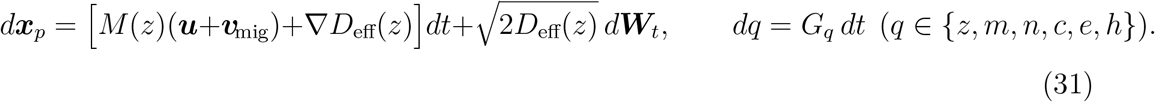

The associated forward Kolmogorov (Fokker–Planck) equation for *f* (***x***, *t*, ***q***), under the reflecting physical wall (Eq. 10), is

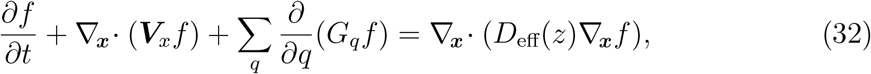

with ***V***_*x*_ = *M* (*z*)(***u*** + ***v***_mig_). There is no separate wall-source term: capture, release and immobilisation are all carried by the internal dynamics of *z* and the capture-dependent transport coefficients, and platelet number is conserved under the reflecting boundary.

**Proposition 2** (Formal Fokker–Planck / Liouville correspondence). *Equation* (32) *is the forward equation of the Itô SDE* (31): *for* 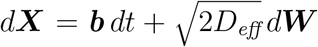 *with* ***b*** = ***V***_*x*_ + ∇*D*_*eff*_, *the diffusion operator ∂*_*i*_*∂*_*j*_(*D*_*eff*_*δ*_*ij*_*f* ) = Δ(*D*_*eff*_*f* ) *combines with the* ∇*D*_*eff*_ *drift to give exactly* ∇_***x***_· (*D*_*eff*_∇_***x***_*f* ). *In the deterministic limit D*_*eff*_ → 0 *it reduces to the Liouville equation, along whose characteristics—the single-platelet ODEs—the density obeys*

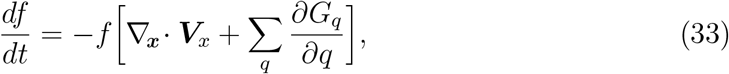

*i*.*e. f J is conserved with J the flow-map Jacobian; f is constant only if the phase-space flow is divergence-free*.

*Proof*. The drift correction ∇*D*_eff_ is the standard Itô-to-Fickian device [28]; this is a formal derivation: *D*_eff_ vanishes where shear vanishes and the wall-zone indicator is non-smooth, so we do not claim global well-posedness. The Liouville relation follows from the material derivative and Liouville’s formula.

**Proposition 3** (Well-posedness of the reduced state). *Under nonnegative rate constants and the stated flux forms, the box* [0, 1]^4^ *for* (*z, m, n, h*) *and the nonnegative cone for* (*c, e*) *are forward-invariant: each bounded variable has zero inward flux at* 0 *and nonpositive rate at* 1, *and J*_*leak*_, *J*_*SERCA*_, *J*_*PM*_ *keep c, e* ≥ 0. *All right-hand sides are locally Lipschitz, giving local existence and uniqueness. We claim boundedness on finite intervals under bounded inputs, but we do* not *assert the a priori bound c* ≤ *c*_*ext*_: *IP*_3_*R release can transiently raise c above c*_*ext*_, *and any global bound must be derived from the specific parameter set rather than assumed*.

Macroscopic observables are moments of *f* : *P* = ∫ *f d****q*** and 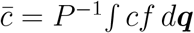.

## 10 Nondimensionalisation and Regime Numbers

With length *L*, velocity *U*, time *T* = *L/U*, and Re = *ρUL/µ*:

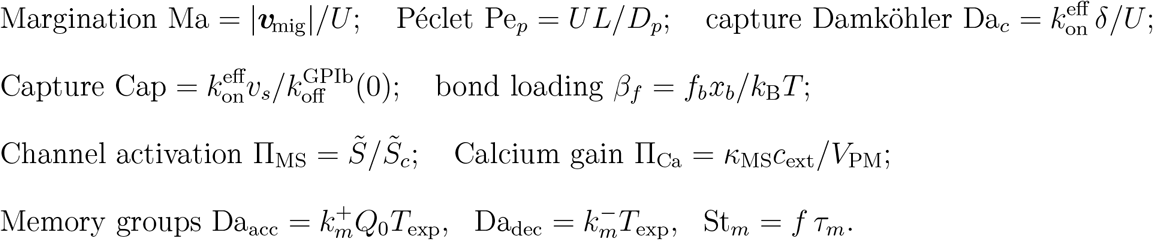

The capture Damköhler number 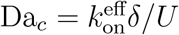 compares the wall-zone residence time *δ/U* to the capture time 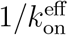 and is dimensionless; it replaces the earlier surface coefficient *k*_capture_, which is no longer part of the model. The memory response is governed by the triple (Da_acc_, Da_dec_, St_*m*_), not by 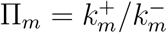 alone; Π_*m*_ is retained only as a coarse saturation index. The time-averaged normalised load is 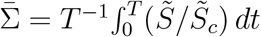.

Figure 2 shows the regime structure. Panel A is the monotone quasi-steady calcium over (effective tension, gating sharpness), separating low-calcium-priming, transitional, and high-mechanosensitive-priming regions (Part I does not model aggregation or thrombus formation, so these are priming regions, not hemostatic/thrombotic states). Panel B is the *absolute* magnitude of the conditioning-dependent conditioning–test signature (Section 12) over (*τ*_*m*_, *λ*_*m*_), using an absolute scale to avoid the small-denominator instability of a ratio metric. The previous “memory-attributable pulsatile advantage” map is withdrawn, because that contrast is a convexity effect rather than memory (Fig. 3D).

**Figure 2:**
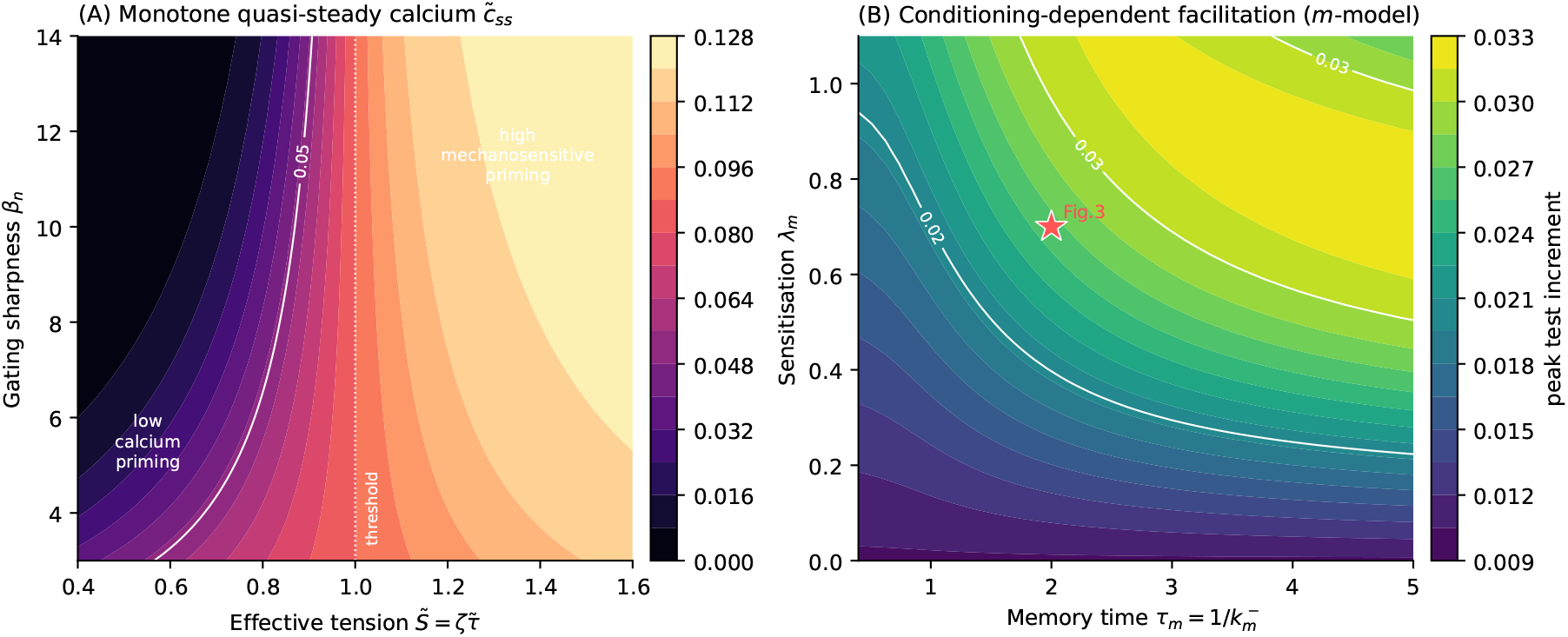
Priming regime map (representative Tier-3 parameters, uncalibrated). **(A)** Monotone quasi-steady cytosolic calcium 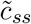 over effective tension 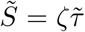 and gating sharpness *β*_*n*_; the dotted line marks the threshold. **(B)** Absolute peak conditioning–test calcium increment (the conditioning-dependent facilitation generated by the *m*-model, Fig. 3) over memory time *τ*_*m*_ and sensitisation *λ*_*m*_; the star marks the operating point of Fig. 3. Absolute scales are used throughout; all values are illustrative outputs of uncalibrated parameters.

**Figure 3:**
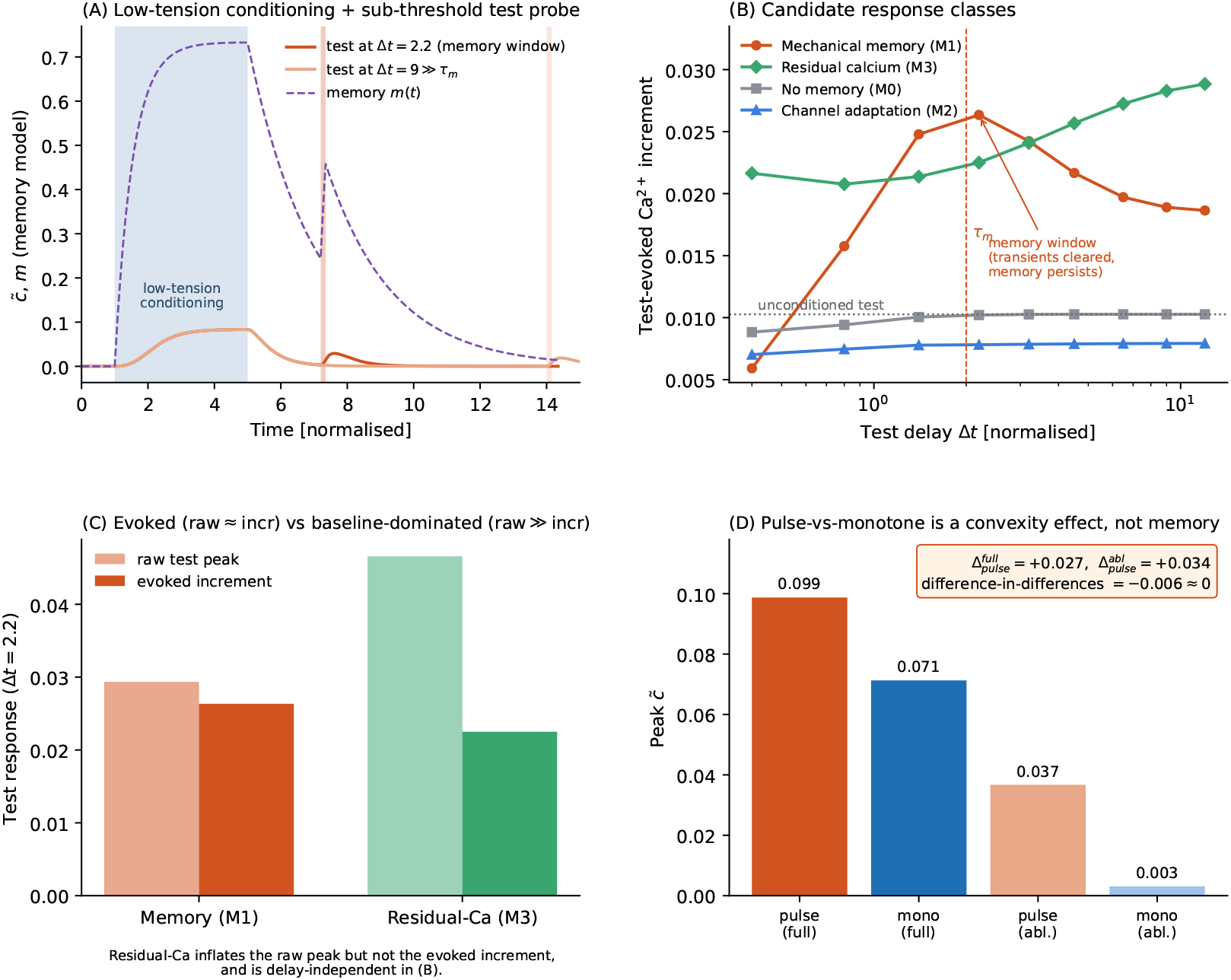
Mechanical memory is identified by the conditioning–test signature, not by the pulsatile-versus-monotone contrast. Normalised system, four competing models with identical downstream calcium readout. **(A)** Protocol: a low-tension conditioning block (charges *m*, minimal calcium) then a fixed sub-threshold test probe; example memory-model traces of 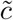 and *m* at a short (Δ*t* = 2.2 ≈ *τ*_*m*_) and long (Δ*t* = 9) delay. **(B)** Test-evoked calcium increment versus delay for the four reduced candidate models; the dotted line marks the unconditioned-test response. Within this comparator set, the memory model (M1) produces a delay-dependent enhancement peaking in the memory window and decaying on *τ*_*m*_; adaptation (M2) is flat/depressed; residual calcium (M3) is elevated but delay-independent. Adhesive- and store-persistence mechanisms were not simulated (see text). **(C)** Raw test peak versus evoked increment at Δ*t* = 2.2: memory is evoked (raw≈increment), whereas residual calcium is baseline-dominated (raw≫increment). **(D)** Four-condition contrast at equal mean shear (pulse/monotone × full/ablated): the difference-in-differences is ≈ 0, confirming that the pulsatile excess is a convexity effect, not memory. All values are illustrative outputs of uncalibrated parameters.

## 11 Calibration Strategy

Parameters are partitioned into three tiers (full listing, units, provenance and uncertainty in Table 2). **Tier 1** (fixed physical constants); **Tier 2** (fitted from published data, Steps A–D); **Tier 3** (memory and load-transfer parameters requiring new waveform-resolved experiments).

### Step A (hemodynamics)

Validate *τ, g*_*τ*_ against particle image velocimetry in stenotic microchannels [13]. **Step B (margination)**. Fit *D*_0_, *χ*_*m*_, 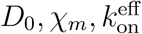 to near-wall concentration profiles versus hematocrit and RBC deformability [6]. **Step C (capture)**. Fit the catch-slip parameters to rolling-velocity distributions and tether lifetimes; the law predicts a non-monotone lifetime versus force testable against single-molecule data [18, 19]. **Step D (mechanotransduction and calcium)**. Fit the gating and calcium parameters to calcium transients under controlled shear [10], with and without GsMTx4. Because GsMTx4 is a gating-modifier rather than a selective pore block [22], the ±GsMTx4 contrast *constrains κ*_MS_ but does not uniquely isolate it.

### Identifiability

Because 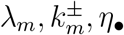 and 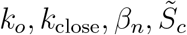 can compensate, calibration must include local and global sensitivity analysis, structural identifiability where feasible, profile likelihoods or Bayesian posteriors, and out-of-sample validation before any numerical output is reported as a prediction.

## 12 Testable Predictions

### 12.1 Prediction 1: conditioning–test facilitation is the memory signature

#### Why the pulsatile-versus-monotone contrast fails

A natural but incorrect test compares pulsatile and monotone shear at equal mean. Because the gating map is a convex sigmoid in the sub-threshold range, intermittent high values produce more response than a constant mean even with *no* memory (Jensen’s inequality). Integrating the reduced system (Fig. 3D) gives, at equal mean, peak calcium of 0.099 (pulsatile, full) versus 0.071 (monotone, full), but also 0.037 (pulsatile, ablated) versus 0.003 (monotone, ablated): the difference-in-differences ( 0.099 − 0.071) − (0.037 − 0.003) ≈ −0.006 ≈ 0. The pulsatile excess is therefore a convexity effect, not memory, and the earlier headline number is withdrawn.

#### The history-sensitive conditioning–test

Memory is identified by a **conditioning– test protocol** (Fig. 3A–C). A *low-tension* conditioning block charges *m* through *Q* while staying below the gating threshold, so it evokes negligible channel/calcium transients and leaves a clean baseline. A fixed *sub-threshold* test pulse then probes the carried-over state at delay Δ*t*. The four competing models make distinct, falsifiable predictions for the test-evoked calcium increment (Fig. 3B):

**M1 mechanical memory:** a delay-dependent enhancement that appears once conditioning transients decay, peaks in the window *τ*_channel_ < Δ*t* < *τ*_*m*_, and *decays on the memory time τ*_*m*_;

**M0 no memory:** flat (the test is always sub-threshold);

**M2 channel adaptation:** flat or depressed, with no facilitation;

**M3 residual calcium:** elevated but *delay-independent* and baseline-dominated—the raw test peak is inflated while the evoked *increment* is not (Fig. 3C), distinguishing it from memory.

Within the four reduced candidate models examined here, the phenomenological memory model produces a distinct delay-dependent facilitation profile, whose primary qualitative signatures are the *delay-dependence* of the increment and its *decay on τ*_*m*_. These are candidate experimental discriminators, not a unique identification of a memory mechanism (see limitation below). The magnitude is a modest, uncalibrated output.

#### Comparators not included

Persistent VWF activation, GPIb–VWF adhesive occupancy *z*, intracellular-store state (*c, e, h*), and other slowly relaxing mechanochemical processes were *not* included as separate comparator models here and may generate overlapping conditioning–test profiles. The protocol is therefore presented as a framework for discriminating candidate mechanisms, not as an assay that uniquely isolates a latent memory state; mechanistic attribution will require simultaneous measurement of calcium together with adhesion or rolling state and the relevant recovery times.

#### Central claim (revised)

*A phenomenological history variable can generate conditioningdependent platelet facilitation whose magnitude decays with the inter-stimulus interval on the memory time τ*_*m*_. *The conditioning–test protocol provides an experimental framework for determining whether an additional sensitisation state is required beyond known adhesive, channel and intracellular-store persistence*.

#### Operational controls

Because GsMTx4 is a gating-modifier, not a memory switch [22], the in-silico ablation *λ*_*m*_ = 0 is *not* experimentally equivalent to GsMTx4: a reduced response under GsMTx4 demonstrates mechanosensitive-channel involvement, not the existence of a memory state. Memory must instead be tested by the interval-dependence in Fig. 3B. A store-content control (thapsigargin-releasable pool) must be matched across conditions—a store-depletion control, not a maximal-IP_3_ stimulus—and ADP scavenging with apyrase excludes paracrine contributions.

### 12.2 Geometry-resolved corroboration under stenotic hemodynamics

The reduced systems of Fig. 3 use idealised normalised waveforms. To check that the waveform behaviour of Eq. (30) is not an artefact of that idealisation, we drove the memory equation with wall-shear-stress histories obtained from geometry-resolved computational fluid dynamics (CFD).

#### Setup and scope

Transient two-dimensional simulations were performed in idealised microchannels containing either a 50% or a 70% area-reduction stenosis (Fig. 4A–B), the geometry class in which stenotic wall-shear amplification [13] and shear-gradient-driven platelet accumulation [7] are characterised. Oscillatory inlet conditions (1 Hz, 50% duty cycle) and matched constant-flow controls were prescribed so that both delivered identical time-averaged flow rates. Wall-shear-stress histories were extracted at the throat (*x* = 0 mm), where forcing is largest. These simulations are Newtonian, rigid-walled and cell-free: they contain no red blood cells and no platelets, and therefore *prescribe the forcing field* for Module VI in isolation rather than exercising Modules II– V. Consistent with Fig. 3, only the shear contribution to the source term in Eq. (30) was retained (*η*_*τ*_ = 1, *η*_*σ*_ = *η*_*z*_ = 0), so that *Q*(*t*) = *τ* (*t*)*/τ*_ref_.

**Figure 4:**
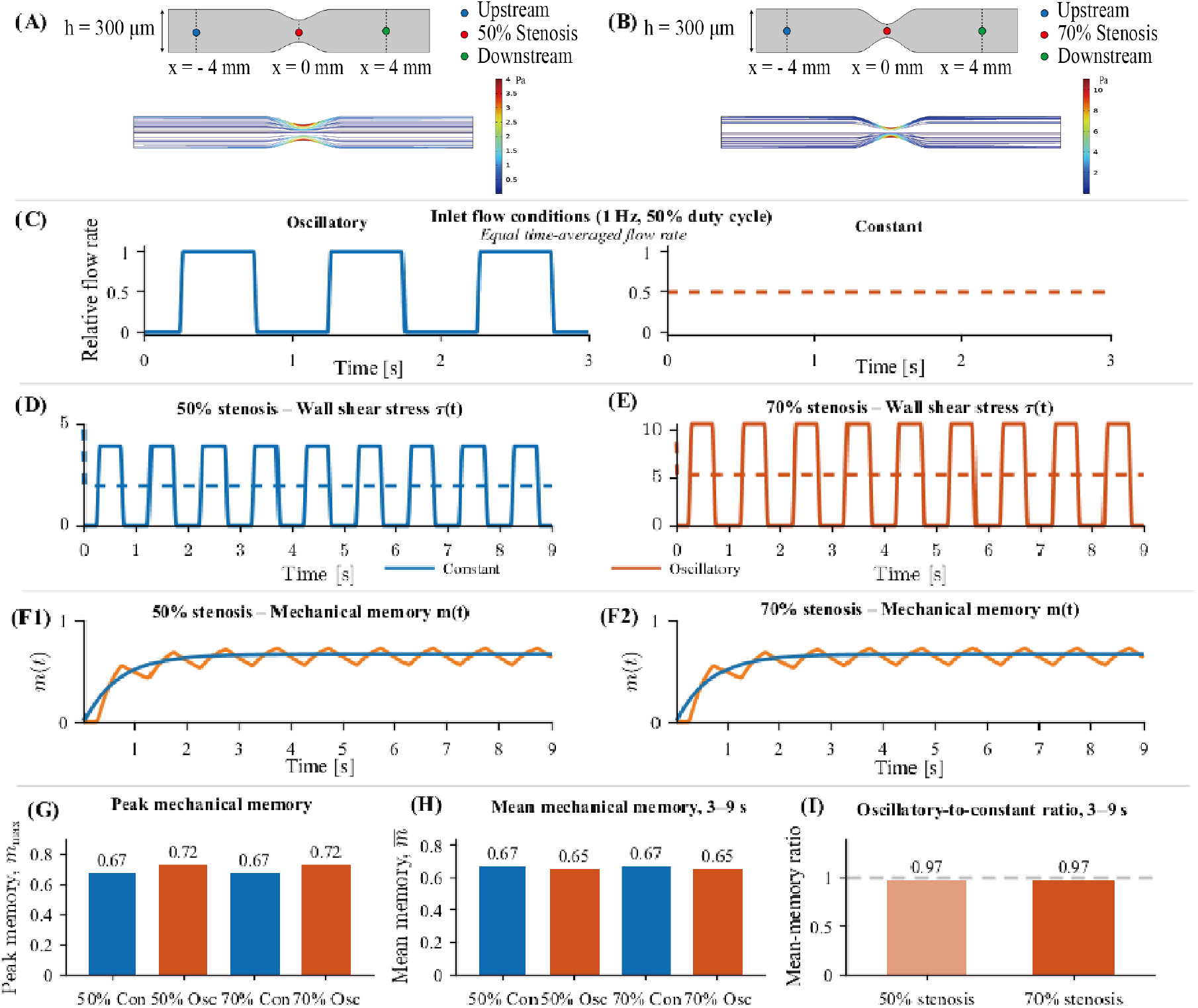
Geometry-resolved CFD forcing of the memory equation under oscillatory and constant shear. **(A–B)** Two-dimensional microchannel geometries with 50% and 70% stenosis and the wall-data extraction locations. **(C)** Oscillatory (1 Hz, 50% duty cycle) and constant inlet conditions with identical time-averaged flow rates. **(D–E)** CFD-derived wall-shear-stress histories at the stenosis throat; increasing stenosis severity produces substantially larger absolute wall shear stress. **(F1–F2)** Mechanical-memory trajectories obtained from the normalised wall-shear-stress histories. Despite the substantial differences in absolute wall shear stress shown in (D–E), the normalised responses from the 50% and 70% stenosis geometries collapse onto nearly identical waveform-dependent trajectories. **(G)** Peak memory. **(H)** Mean memory over 3–9 s. **(I)** Oscillatory-to-constant mean-memory ratio. Oscillation increases the transient *peak* memory response, an expected consequence of *m* partially tracking the instantaneous forcing *τ* (*t*) and not by itself evidence of a memory effect, while leaving the *mean* memory essentially unchanged (ratio ≈ 0.97), consistent with 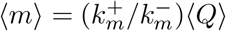. Simulations are Newtonian, rigid-walled, and cell-free; parameters are representative and uncalibrated.

#### Flow fields and the normalization

The flow fields showed the expected acceleration through the throat, with progressively stronger velocity focusing and wall-shear amplification as stenosis severity increased; the 70% geometry generated substantially larger absolute wall shear stresses than the 50% geometry (Fig. 4D–E,), consistent with the severity dependence reported for stenotic flows [13]. Oscillatory flow produced periodic high- and low-shear phases, whereas the constant-flow controls were time-invariant at the mean wall shear stress of the corresponding oscillatory condition (Fig. 4C–E). Because the object here is the contribution of temporal shear-history structure rather than of absolute magnitude, each history was normalised by the peak wall shear stress of the corresponding oscillatory case before evaluation of Eq. (30). The two severities then serve as an internal consistency test: if geometry-dependent magnitude effects are removed by the normalisation, the memory responses should collapse.

#### Results

They do. The normalised trajectories from the 50% and 70% geometries converge to nearly identical responses (Fig. 4F1–F2) despite the large difference in absolute wall shear stress, confirming that the normalisation isolates waveform history. Under constant forcing the memory variable approaches the analytical steady state 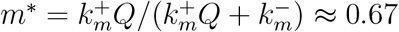. Under oscillatory forcing, high-shear phases produce transient excursions to *m*_max_ ≈ 0.72, an increase of approximately 8% in *peak* memory in both geometries (Fig. 4G). Over the interval 3–9 s, however, the *mean* memory level is comparable between conditions and marginally lower under oscillation, giving an oscillatory-to-constant mean-memory ratio of approximately 0.97 (Fig. 4H–I).

#### Interpretation, and what this does not show

The peak excursion is not evidence of memory: with *η*_*τ*_ = 1 the state *m* partially tracks the instantaneous *τ* (*t*), so a waveform with a higher peak yields a higher peak *m* whether or not any sensitisation follows. The informative quantity is the mean, and the ratio of 0.97 ≈ 1 is a geometry-resolved confirmation of the linearised result 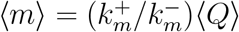 (Section 8): equal-mean monotone and pulsatile forcing produce essentially equal mean memory even under realistic stenotic histories, not only under idealised square waves. This independently supports the withdrawal of the pulsatile-versus-monotone claim in Section 12 above. These runs include no ablation arm (*λ*_*m*_ = 0) and no downstream calcium readout, so they cannot by themselves discriminate memory from convexity; that role remains with the difference-in-differences and conditioning–test contrasts of Fig. 3B,D. What the CFD adds is threefold: the normalisation collapse across severities; the demonstration that mean memory is waveform-insensitive under geometry-resolved forcing; and the dissociation between peak and mean memory, which is precisely why a protocol probing *transient carry-over between discrete stimuli* rather than a mean-waveform contrast is the appropriate assay for the history state.

### 12.3 Prediction 2: RBC stiffness reduces delivery and ensemble calcium

The model directly supports the chain *D*_RBC_ ↓⇒ ***v***_mig_ ↓⇒ near-wall enrichment↓⇒ *J*_enc_ ↓⇒ capture events↓ (captured count ∝ *zN*_tot_)⇒ total surface-associated calcium signal↓. Accordingly, in whole blood at 40% hematocrit and mean wall shear rate 500 s^−1^, stiffening RBCs by 0.01% glutaraldehyde fixation should reduce (a) near-wall platelet enrichment, (b) entry into the wall-interaction zone *J*_enc_, (c) GPIb-mediated capture-event frequency and captured platelet number, and hence (d) total surface-associated (ensemble) calcium fluorescence, relative to unfixed blood [6].

*The model does not currently predict a reduction in calcium amplitude per captured platelet*: once a platelet has bound, its calcium response depends on local shear, bond state and transmitted load, which are not explicitly coupled to RBC deformability except through the collision term. Per-captured-platelet calcium is therefore an experimental *discriminator* rather than a prescribed model outcome: an unchanged per-platelet amplitude would support a predominantly transport-mediated effect, whereas a reduction would indicate an additional RBC-mechanics-to-load coupling not represented in Part I. Reporting both the ensemble signal and the per-platelet amplitude is therefore diagnostic. Glutaraldehyde stiffening additionally requires controls for altered blood viscosity, RBC aggregation, RBC surface chemistry, hemolysis, nonspecific platelet–RBC adhesion, platelet count, and plasma VWF concentration. Aggregate surface coverage, which requires platelet–platelet aggregation, is not a Part I state variable and is deferred to Part II.

### 12.4 End-to-end illustration

Figure 5 integrates the assembled modules for a single platelet: margination delivers it to the wall, GPIb*α* capture progressively immobilises it, the transmitted load crosses the gating threshold, memory accumulates, channels open, and a localised calcium transient results-demonstrating that the coupled framework produces the intended delivery-to-calcium chain. Thus, a reduced, representative simulation, not a pre-diction.

**Figure 5:**
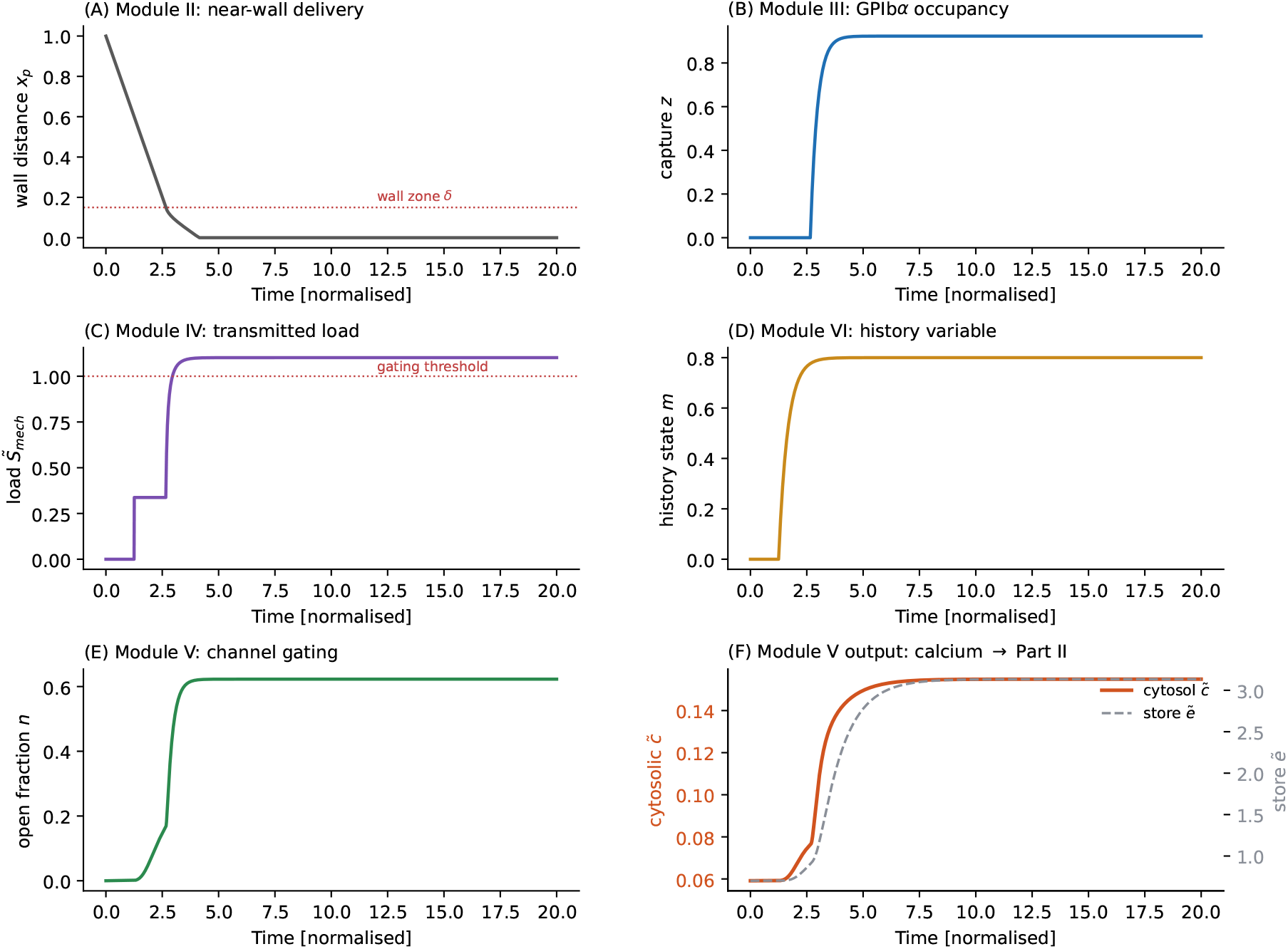
End-to-end reduced single-platelet simulation. A single deterministic integration of the coupled modules, initialised at the resting steady state (*c*_0_ = 0.059, *e*_0_ = 0.702 normalised): **(A)** near-wall delivery (wall distance *x*_*p*_ falling into the wall zone); (**B**) GPIb*α* occupancy *z* (progressive immobilisation, not exact arrest); **(C)** transmitted load 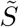 crossing the gating threshold; **(D)** history state *m*; **(E)** channel open fraction *n*; **(F)** cytosolic (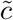, left axis) and intracellular-store (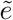, right axis) calcium on *separate* scales, the Module-V output feeding Part II. The calcium mass-balance identity holds to machine precision (residual ∼10^−16^; see Reproducibility). Normalised; representative, uncalibrated parameters.

## 13 Discussion

### 13.1 Position relative to existing frameworks

The present study does not claim to establish a unique molecular mechanism of platelet history dependence. Instead, it embeds a candidate phenomenological history state within a unified delivery-to-calcium framework and identifies measurements capable of distinguishing this state from other persistent mechanochemical processes. Fluid-mechanical models resolve hemodynamics but threshold activation [13]; coagulation models prescribe activation [14]; cell-resolved models omit intracellular state [15]. We do not claim to be the first unified theory: the contribution is the explicit, dimensionally consistent *chain to intracellular calcium*, together with a history-dependent priming hypothesis framed as a *competing model* against channel adaptation and residual-calcium priming rather than asserted. The four elements we advance are the mechanical-memory state *m*(*t*); the VWF decomposition with an objective extensional measure and bounded kernel; a two-pathway catch-slip law parameterisable from single-molecule data; and a single-load membrane stimulus coupled to a dimensionally consistent cytosol–store calcium model with extracellular influx.

### 13.2 Consistency with the literature

Table 1 records the correspondence between ten studies and the modules, using neutral categories that state the *kind* of support. Modules I–V are each supported at the level of mechanism or parameterisation. Module VI (memory) has the weakest direct precedent: history dependence is documented in bulk/ensemble assays [1, 3] but not at the single-platelet, waveform-resolved level. Precisely because direct evidence is absent, the conditioning–test discriminator (Fig. 3) constitutes a genuine test rather than a post-hoc rationalisation. We caution against describing the framework as validated: Tier-3 calibration and confirmation of Predictions 1–2 are prerequisites.

**Table 1:** Correspondence between Part I elements and published literature. Categories: *Potential calibration dataset* (study could directly set the relevant equation once fitted); *Mechanism* (qualitative mechanism supported); *Trend* (population/aggregate trend only). No parameter fit has yet been performed; the framework is trend-consistent, not validated.

| No. | Study | Module | Key correspondence | Category |
| --- | --- | --- | --- | --- |
| 1 | Hellums (1994) [1] | I | Shear stress is a direct agonist above a threshold. | Mechanism |
| 2 | Bark & Ku (2010) [13] | I | Severe stenoses give order-of-magnitude higher wall shear. | Trend |
| 3 | Nesbitt et al. (2009) [7] | I,III | Shear gradients drive localised thrombus growth. | Mechanism |
| 4 | Czaja et al. (2020) [6] | II | Stiffened RBCs reduce near-wall enrichment. | Potential calibration dataset |
| 5 | Schneider et al. (2007) [5] | III | Extensional flow unfolds and activates VWF. | Mechanism |
| 6 | Ruggeri et al. (2006) [2] | III | Activation-independent adhesion via immobilised VWF. | Mechanism |
| 7 | Yago et al. (2008) [18] | III | GPIbα-A1 catch bonds (lifetime maximum vs force). | Potential calibration dataset |
| 8 | Liu et al. (2022) [21] | III | Rapid SIPA of nonactivated platelets via VWF-GPIb-A1. | Mechanism |
| 9 | Ilkan et al. (2017) [10] | V | Shear-mediated Ca<sup>2+</sup> entry reduced by GsMTx4/gadolinium. | Potential calibration dataset |
| 10 | Zhao et al. (2021) [11] | V | Piezo1 gain-of-function accelerates thrombosis. | Mechanism |

### 13.3 Assumptions and limitations

**(A)** Newtonian blood (large-vessel stenotic flows). **(B)** Overdamped platelets. **(C)** *S*_mech_ is an effective mechanosensory load, not a bilayer tension. **(D)** Lumped channel population. **(E)** Prescribed IP_3_ in Part I. Principal limitations: one-way fluid–platelet coupling; no coagulation/fibrin; no endothelial biology; no shape-change; no paracrine ADP loop (Part II); and memory is not yet distinguished experimentally from adaptation or residual-calcium priming—the model-selection question Fig. 3 is designed to resolve. The geometry-resolved simulations of Section 12.2 are additionally two-dimensional, rigid-walled and cell-free, and are not validated against velocimetry: they supply a realistic forcing history to Module VI rather than a validated hemodynamic prediction, and they neither exercise Modules II–V nor discriminate between the competing models. The regime boundaries and magnitudes in Figs. 2–5 are representative, uncalibrated outputs; only the qualitative topology and the delay-dependent memory signature are claimed to be robust.

## 14 Conclusions

We have presented a reduced, six-module mechanobiological framework linking vessel-scale hemodynamic forcing to mechanosensitive calcium entry through dimensionally consistent couplings. Its central prediction is reformulated around a history-sensitive conditioning–test signature, and we show explicitly that the previously proposed pulsatile-versus-monotone contrast is a convexity effect rather than a test of memory. The single-platelet stochastic dynamics and the population balance form a Fokker– Planck pair, with spatially varying diffusivity handled by an explicit drift correction. The framework is offered as a calibratable, hypothesis-generating theory—not a validated model—whose Tier-3 parameters and history hypothesis are the targets of the experiments proposed here. The principal contribution is therefore not proof of a distinct memory molecule or pathway, but a coherent mathematical framework and an experimentally actionable strategy for determining which persistent states govern history-dependent platelet priming. The cytosolic calcium *c*(*t*) is the biochemical input to Part II.

### Reproducibility

The Python scripts generating Figs. 2, 3, 4 and 5 (regime map, conditioning–test discriminator with the four candidate models, CFD-driven memory trajectories, and the end-to-end single-platelet simulation) are provided with the submission, together with the complete parameter set (Table 2), initial conditions, reference scales, conditioning and test waveform definitions, and normalisation conventions. The general framework contains stochastic transport (Eq. 31); the reduced simulations use deterministic representative trajectories, so no random seed is required. Integration is fixed-step (forward Euler, step *h* = 10^−3^ normalised). Under successive time-step refinement (*h, h/*2, *h/*4) the principal outputs—peak cytosolic calcium and the conditioning-induced increment—change by less than 0.2%, i.e. below the 1% tolerance. For the cytosol–store model the calcium mass-balance identity 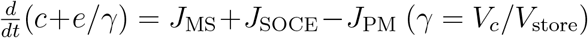 holds to machine precision (maximum residual ∼ 10^−16^), confirming that the internal fluxes redistribute rather than create calcium. No calibration to experimental data is claimed.

**Table 2:** Complete parameter set with units, provenance and status. Tier 1: fixed. Tier 2: fittable from cited data. Tier 3: requires new waveform-resolved experiments. Normalised figure values are representative and uncalibrated.

| Parameter | Meaning | Units | Tier / provenance |
| --- | --- | --- | --- |
| $R_p$ | platelet radius | $\mu\text{m}$ ($\approx 1.5$) | 1 / morphometry |
| $a_{\text{RBC}}$ | RBC radius | $\mu\text{m}$ ($\approx 4$) | 1 / morphometry |
| $\mu$ | plasma viscosity | $\text{mPa s}$ ($\approx 3.5$) | 1 / rheology |
| $k_{\text{B}}T$ | thermal energy | $4.11 \times 10^{-21} \text{ J}$ | 1 / physical |
| $c_{\text{ext}}$ | extracellular $\text{Ca}^{2+}$ | $\text{mM}$ ($\approx 1.8$) | 1 / physiology |
| $V_c/V_{\text{ER}}$ | cytosol:ER volume | $\approx 10$ | 1 / [24] |
| $A_{\text{con}}$ | contact area | $\mu\text{m}^2$ (0.2–1) | 1–2 / geometry |
| $D_0$ | diffusivity prefactor | — (0.025–0.10) | 2 / [6, 16] |
| $\chi_m$ | margination mobility | $\text{m s}^{-1}$ | 2 / [6] |
| $\delta$ | wall-interaction zone width | $\mu\text{m}$ ($\sim R_p$) | 1–2 / geometry |
| $\tau_v, \beta_\tau$ | VWF threshold, sharpness | $\text{Pa}, \text{Pa}^{-1}$ | 2 / [5] |
| $A_g, K_g$ | gradient gain, half-sat | —, $\text{Pa m}^{-1}$ | 2 / [7] |
| $A_\epsilon, K_\epsilon$ | extensional gain, half-sat | —, $\text{s}^{-1}$ | 2 / [5] |
| $k_{\text{on}}^{\text{eff}}$ | effective on-rate | $\text{s}^{-1}$ | 2 / [18] |
| $k_{\text{off}}^{A,B}$ | slip/catch off-rates | $\text{s}^{-1}$ | 2 / [18, 19] |
| $x_b^{A,B}$ | slip/catch distances | nm | 2 / [18] |
| $N_{\text{ref}}$ | rolling→immobilisation scale | bonds | 2 / [21] |
| $\kappa_{\text{MS}}$ | mechanosensitive rate | $\text{s}^{-1}$ | 2 / [10] |
| $V_{\text{SERCA}}, K_{\text{SERCA}}$ | SERCA max flux, half-sat | $\mu\text{M s}^{-1}, \mu\text{M}$ | 2 / [24] |
| $V_{\text{PM}}, K_{\text{PM}}$ | PMCA/NCX max flux, half-sat | $\mu\text{M s}^{-1}, \mu\text{M}$ | 2 / [27] |
| $V_{\text{SOCE}}, K_{\text{SOCE}}, r$ | SOCE max flux, half-sat, Hill | $\mu\text{M s}^{-1}, \mu\text{M}, \text{—}$ | 2 / [27] |
| $k_{\text{IP3R}}, K_{\text{IP3}}, K_{\text{act}}, K_{\text{inh}}$ | IP<sub>3</sub>R kinetics | $\text{s}^{-1}, \mu\text{M}$ | 2 / [24, 25] |
| $k_{\text{leak}}$ | store leak | $\text{s}^{-1}$ | 2 / [24] |
| $k_o, k_{\text{close}}, \beta_n, \tilde{S}_c$ | gating kinetics, sharpness, threshold | $\text{s}^{-1}, \text{—}, \text{—}$ | 2–3 / [9, 10] |
| $k_i, k_r$ | adaptation in/recovery (M2) | $\text{s}^{-1}$ | 2–3 / [23] |
| $\mathcal{T}_0$ | baseline load transfer | — | 3 / new |
| $\lambda_m$ | memory sensitisation | — | 3 / new |
| $k_m^+, k_m^-$ | memory accumulation/decay | $\text{s}^{-1}$ | 3 / new |
| $\eta_\tau, \eta_\sigma, \eta_z$ | memory source weights | — | 3 / new |
| [IP<sub>3</sub>] | agonist-set IP<sub>3</sub> (Part I input) | $\mu\text{M}$ | input / [25] |

The geometry-resolved simulations of Fig. 4 solve the incompressible Navier–Stokes equations in two dimensions for a Newtonian fluid (*µ* = 3.5 mPa s, *ρ* = 1060 kg m^−3^) in rigid-walled channels with a 50% or 70% area-reduction stenosis, with no-slip walls, a prescribed inlet flow rate (oscillatory: 1 Hz, 50% duty cycle; constant: matched time-averaged flow rate) and a zero-pressure outlet. [Solver and version, channel dimensions and stenosis profile, throat Reynolds and Womersley numbers, mesh size with near-wall refinement, time-step, and the mesh- and time-step-refinement study demonstrating that throat wall shear stress changes by less than the stated tolerance under successive refinement, to be inserted.] Extracted wall-shear-stress histories at the throat, the normalisation reference *τ*_ref_, and the script integrating Eq. (30) are provided with the submission. These simulations contain no cells and are not validated against experimental velocimetry; they are used solely to supply a realistic forcing history to Module VI.

## Acknowledgements

[To be added.] **Conflicts of interest:** The authors declare none.

## Notes

### Competing Interest Statement

The authors have declared no competing interest.

